# Epigenomic and 3D genome architecture in naïve and primed human embryonic stem cell states

**DOI:** 10.1101/181123

**Authors:** Stephanie L. Battle, Naresh Doni Jayavelu, Robert N. Azad, Jennifer Hesson, Faria N. Ahmed, Joseph A. Zoller, Julie Mathieu, Hannele Ruohola-Baker, Carol B. Ware, R. David Hawkins

## Abstract

During mammalian embryogenesis changes in morphology and gene expression are concurrent with epigenomic reprogramming. Using human embryonic stem cells representing the pre-implantation blastocyst (naïve) and post-implantation epiblast (primed), our data demonstrate that a substantial portion of known human enhancers are pre-marked by H3K4me1 in naïve cells, providing an enhanced open chromatin state in naïve pluripotency. The naïve enhancer repertoire occupies nine percent of the genome, three times that of primed cells, and can exist in broad chromatin domains over fifty kilobases. Enhancer chromatin states are largely poised. Seventy-seven percent of naïve enhancers are decommissioned in a stepwise manner as cells become primed. While primed topological associated domains are unaltered upon differentiation, naïve domains expand across primed boundaries, impacting three dimensional genome architecture. Differential topological associated domain edges coincide with naïve H3K4me1 enrichment. Our results suggest that naïve-derived cells have a chromatin landscape reflective of early embryogenesis.

## INTRODUCTION

Dynamic changes in the epigenome are concerted with morphological and gene expression changes during early embryogenesis. Soon after fertilization DNA methylation is actively removed from the paternal genome, passively lost from the maternal genome and regained in the post-implantation epiblast(Guo et al., 2014). In addition to resetting the DNA methylome, the early embryonic epigenome maintains an open chromatin structure as repressive heterochromatin is gained later over the course of development, lineage commitment and differentiation(Ahmed et al., 2010; Liu et al., 2004; Sarmento et al., 2004). These changes in histone modifications correlate with the hypothesis that a more open chromatin structure is a key aspect of pluripotency and allows embryonic cells to respond to a broad array of developmental signaling cues(Hawkins et al., 2010; Meshorer et al., 2006).

Pre- and post-implantation pluripotent ESCs provide a system to model epigenomic reprogramming during early embryogenesis and to study changes in pluripotency. Mouse ESCs (mESCs) are currently the primary model for studying mammalian pre-implantation embryos and deemed naïve(Silva and Smith, 2008), while mouse epiblast stem cells (EpiSCs) model the post-implantation embryo and exist in the primed state of pluripotency(Brons et al., 2007; Tesar et al., 2007). Due to a number of similarities between mouse EpiSCs and human ESCs (hESCs), it is now accepted that hESCs exist in the primed state(Nichols and Smith, 2009). However, several groups described the first set of naïve hESCs, where primed hESCs or human iPSCs were induced, or reset, to the naïve state(Chan et al., 2013; Gafni et al., 2013; Hanna et al., 2010; Takashima et al., 2014; Theunissen et al., 2014; Valamehr et al., 2014; Ware et al., 2014). Additionally, new hESC lines were derived, each under a different naïve growth condition(Gafni et al., 2013; Guo et al., 2016; Theunissen et al., 2014; Ware et al., 2014) (for review see(Ware, 2016)). Similar to mouse, naïve hESCs exhibit DNA hypomethylation and two active X chromosomes(Gafni et al., 2013; Theunissen et al., 2016; Ware et al., 2014), hallmarks of the pre-implantation state.

Given the differences between early human and mouse embryogenesis(Blakeley et al., 2015; Rossant, 2015), naïve-derived hESC lines provide an opportunity to study changes that are reflective of early human development and pluripotency. To better our understanding of epigenomic reprogramming as hESCs transition from the pre-implantation to post-implantation state, we present data from whole transcriptome RNA-seq, ChIP-seq for five histone modifications, and topological associated domains (TADs) from in situ DNaseI Hi-C for the naïve-derived Elf1 line(Ware et al., 2014) grown in 2i + Lif + IGF1 + FGF (2iLIF). We include data from cells transitioning from the naïve state (Activin + FGF noted as AF) and compared our results to data from primed H1 hESC(Dixon et al., 2012; Hawkins et al., 2010). Extensive chromatin remodeling occurs at promoters and enhancer elements as cells transition from naïve to primed. Our analysis reveals that naïve hESCs have a more open chromatin structure due to large expansions of H3K4me1 and H3K27ac in the genome. Seventy-seven percent of naïve enhancers are decommissioned in the primed state. TADs are largely stable between pluripotent states, but our data reveal limited naïve specific shifts in TAD boundaries. Overall, these data provide an extensive view of the epigenome and 3D genome for hESC states and a model of epigenomic reprogramming during early human embryogenesis.

## RESULTS

### Gene Expression in Naïve hESCs

Naïve and primed hESCs are expected to have distinct expression profiles, and naïve cells should reflect aspects of human blastocyst gene expression. We performed strand-specific, whole transcriptome RNA-seq in replicate on Elf1 naïve (2iLIF), Elf1 transitioning (AF) and H1 primed (mTeSR) cells of equal cell numbers (Supplementary Figure 1a-c; see methods for growth conditions). We identified differentially expressed genes (DEGs) in a pairwise manner (Fig. 1a, b). The largest number of DEGs was observed between naïve and primed hESCs (Fig. 1b and Supplementary Table 1), signifying just how distinct these cellular states are. Highlighted are several genes known to be upregulated in the human pre-implantation epiblast(Blakeley et al., 2015; Yan et al., 2013) and other genes of interest, indicating that the characteristics we observe for 2iLIF naïve cells are reflective of pre-implantation development.

**Figure 1.**
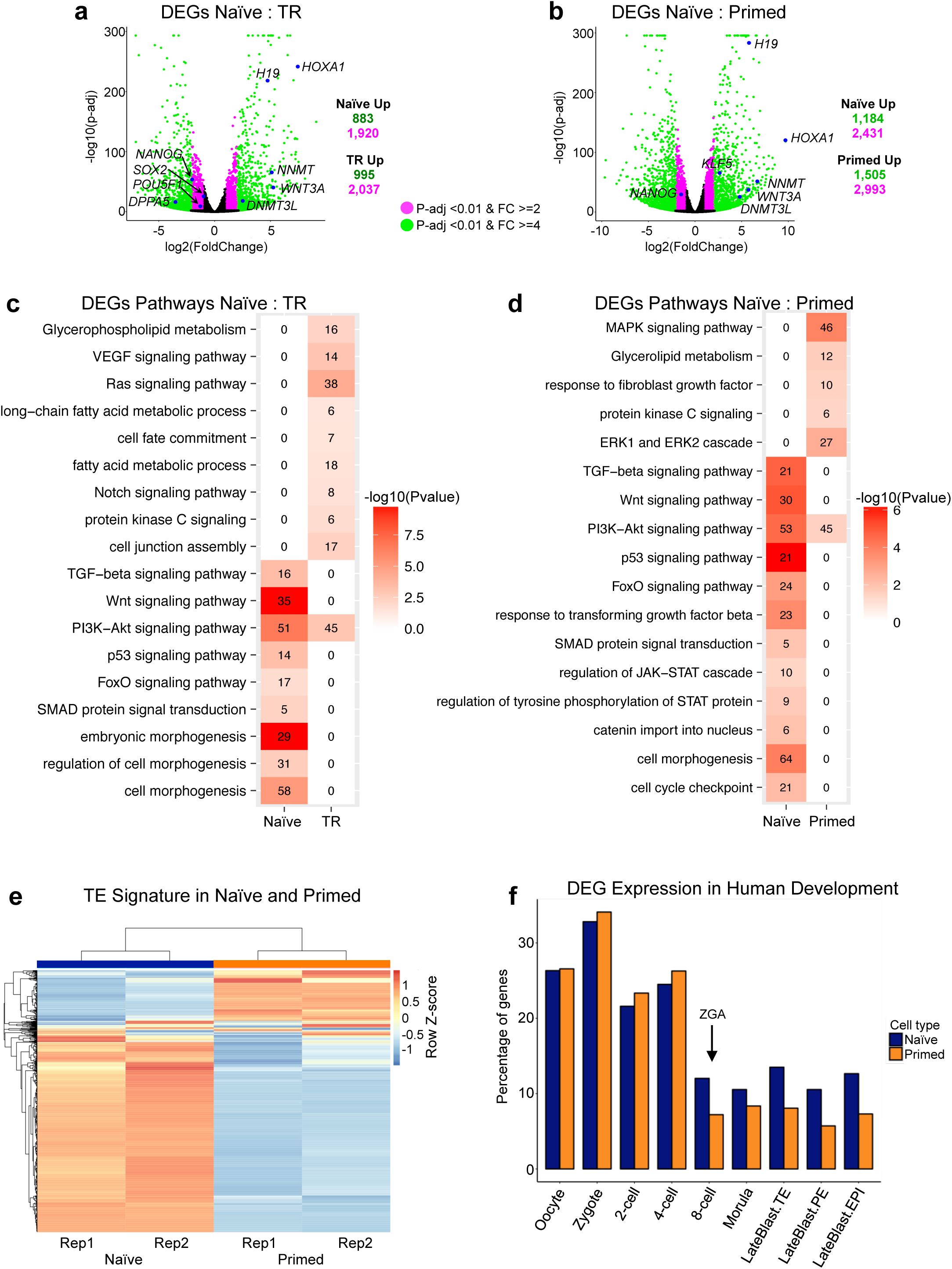
Differential Gene Expression. (a-b) Volcano plot of differentially expressed genes (DEGs) in naïve versus transitioning (a) and naïve versus primed (b) pairwise comparison. Genes in magenta have P-adj < 0.01 and fold change ≥ 2 while genes in green have P-adj < 0.01 and fold change ≥ 4. (c-d) Heatmap showing significantly overrepresented GO terms and KEGG pathways based on DEGs in naïve versus transitioning (c) and naïve versus primed (d) pairwise comparison. (e) Hierarchical clustering of transposable elements gene expression separates naïve from primed hESCs. (f) Percentage of genes upregulated in pairwise comparison of naïve or primed hESCs that are also found to be upregulated in human embryo developmental stages.

We determined gene ontology (GO) categories and KEGG pathways for naïve DEGs, which were significantly enriched for embryo development and pluripotency signaling pathways along with other pathways important during pre-implantation development (Fig. 1c,d). In particular, genes in the TGF-beta pathway were found to be upregulated in naïve cells, including *LEFTY1*, *SMAD3* and *NODAL* (Supplementary Fig. 1d). The TGF-beta pathway was shown to be important for maintenance of *NANOG* in the human epiblast, whereas inhibition of this pathway has insignificant effects on mouse embryos(Blakeley et al., 2015). PI3K-AKT signaling pathway was also enriched, and is known to promote ESC self-renewal through inhibition of ERK signaling pathway(Supplementary Fig. 1e) (Chen et al., 2012). The WNT signaling pathway was enriched for naïve upregulated genes including *WNT8A*, *WNT5B* and *TCF7* (Supplementary Fig. 1f)(Sperber et al., 2015). A number of terms associated with embryonic development and morphogenesis were enriched for naïve upregulated genes. This may foreshadow what happens to the cells of the blastocyst as they prepare to become the embryonic disk of the epiblast.

We identified cell type-specific genes in the different hESC stages by applying a cutoff of a RPKM value greater than or equal to two in one cell type and less than one in the other two cell types (Supplemental Fig 2). Using this cutoff we determined 429 naïve-specific genes, 229 transition-specific genes and 333 primed-specific genes. Compared to the primed states, naïve-specific genes were enriched for GO terms associated with morphogenesis and pattern specification (Supplemental Fig 2). This is due, in part, to the many *HOX* genes that are uniquely expressed in naïve hESCs and not in transitioning or primed cells. Primed cells were enriched for terms associated with extracellular communication and protein/histone demethylation.

**Figure 2.**
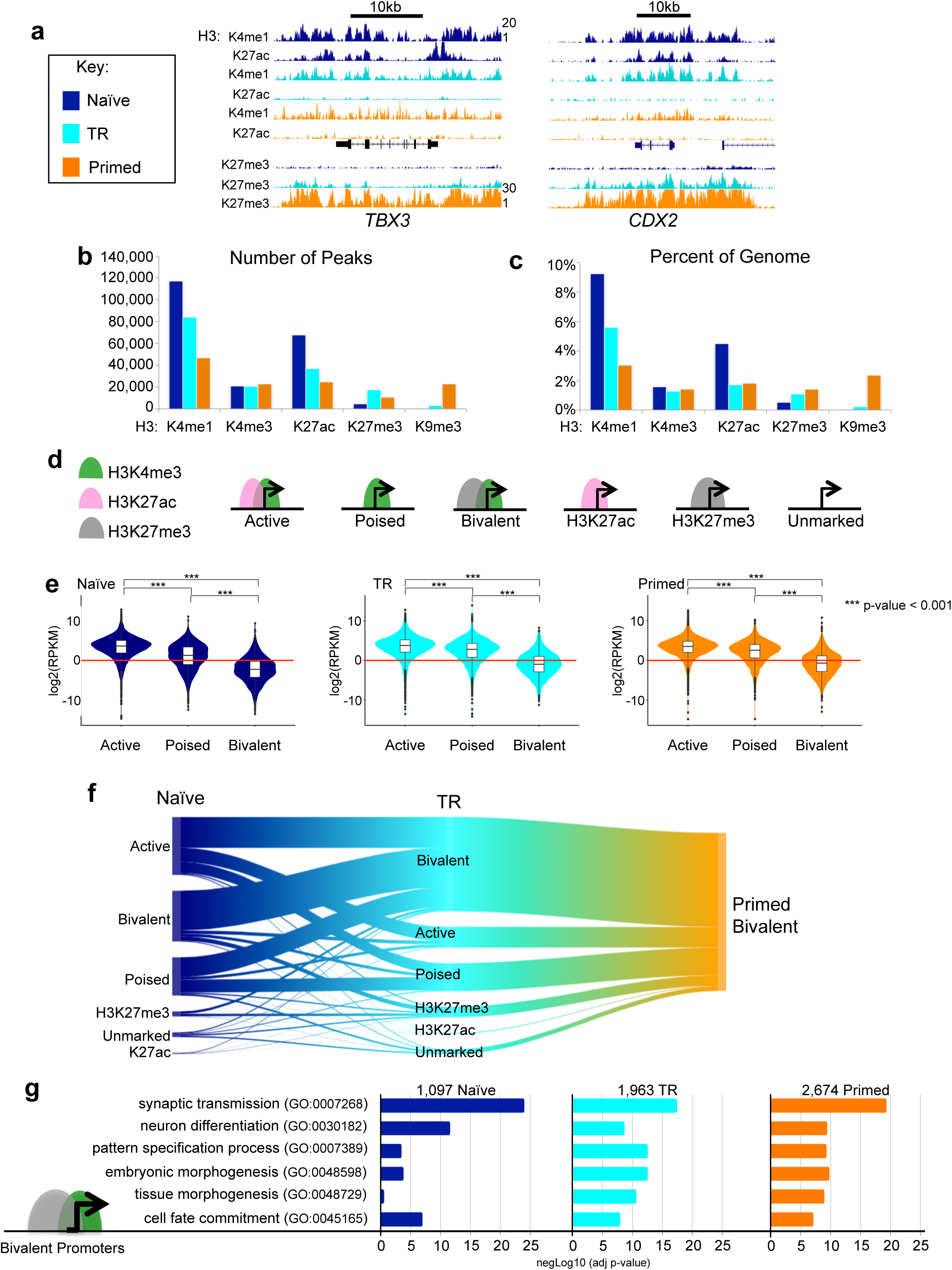
Overview of Chromatin States. Global view of chromatin structure for naïve (navy), transitioning (cyan) and primed (orange) hESCs. These colors are used throughout all figures. (a) UCSC Genome Browser images of *TBX3* and CDX2 gene loci showing enrichment of H3K4mel (RPKM range l-20), H3K27ac (RPKM range 1-20), and H3K27me3 (RPKM range 1-30) in naïve, transitioning and primed cells. (b) The number of ChIP-seq peaks called by MACS with FDR cutoff ≤ 0.05. (c) The percent of genome covered by each histone modification (number of bases divided by effective genorne size: 2.7e+09). (d) Promoters were classified based on the criteria pictured - also see main text. (e) Violin plots showing the distribution of RPKM values of nearest neighboring genes of active, poised and bivalent promoter peaks in each cell type. P-values for pairwise comparisons are computed using two tailed t-tests with pooled SD. P-values are adjusted with Benjamini-Hochberg method. *** P-value < 0.001;. (f) Sankey plot of primed bivalent gene promoters and their origins from the naïve state. (g) Significance of GO Terms from bivalently mark ed gene promoters.

A recent report showed that the transposable element (TE) transcriptome can be used as a state-specific signature in hESCs(Theunissen et al., 2016). Naïve and primed hESCs segregate when clustered on the top 1,000 highly expressed TEs (Fig. 1e). Lastly, we compared upregulated genes to human embryo RNA-seq data from Yan et al.(Yan et al., 2013). We find that a similar percentage of upregulated genes from naïve and primed are expressed in pre-zygotic genome activation stages, while naïve hESCs share more upregulated genes with the post-ZGA embryo than primed (Fig. 1f). This strengthens reports that naïve cells are a good representative model of the pre-implantation stage of human development(Sperber et al., 2015; Ware et al., 2014).

### Global Chromatin Features of Naïve hESCs

To assess global chromatin dynamics between the cellular states, we performed ChIP-seq on five histone modifications from naïve and transitioning cells (Supplementary Table 2,3), and used data previously generated in H1 hESCs for the primed state(Hawkins et al., 2010). These modifications include: H3K4me3 for Pol II-bound promoters(Barski et al., 2007; Guenther et al., 2007; Heintzman et al., 2007), H3K4me1 for enhancers(Heintzman et al., 2009; Heintzman et al., 2007), H3K27ac for active regions(Hawkins et al., 2011; Heintzman et al., 2009), H3K27me3 for Polycomb repressed regions(Bernstein et al., 2006; Boyer et al., 2006), and H3K9me3 for heterochromatin(Bannister et al., 2001; Barski et al., 2007). All five modifications along with ChIP inputs were sequenced in duplicates for both Elf1 naïve and Elf1 transitioning cells for a total of > 270 million and >213 million sequencing reads respectively (Supplementary Table 2,3).

We inspected genes with known expression differences during early embryogenesis through the blastocyst/epiblast stage to ensure our chromatin maps reflect changes during differentiation from naïve to primed. *TBX3* was shown to be expressed in naïve ESCs and human epiblasts(Blakeley et al., 2015). The *TBX3* locus exhibits high levels of H3K4me1 and H3K27ac in naïve hESCs, a reduction of H3K27ac in the transitioning state, followed by a reduction of H3K4me1 and a gain of H3K27me3 in primed hESCs (Fig. 2a). *KLF2*, which was shown not to be expressed in human naïve cells(Blakeley et al., 2015), lacks the H3K27ac modification in all three hESC stages (Supplementary Figure 3a). *CDX2* has active histone modifications in naïve hESCs but transitions to lost acetylation and gained H3K27me3 in primed hESCs (Fig. 2b). CDX2 has been shown to be expressed after blastocyst formation in human embryos and overlaps OCT4 expression in preimplantation embryos(Niakan and Eggan, 2013). Expansion of H3K27me3 domains are also shown at the *HOXA* locus as hESC move from naïve to primed (Supplementary Figure 3b). Next, we asked whether these trends observed at specific loci held true genome-wide.

**Figure 3.**
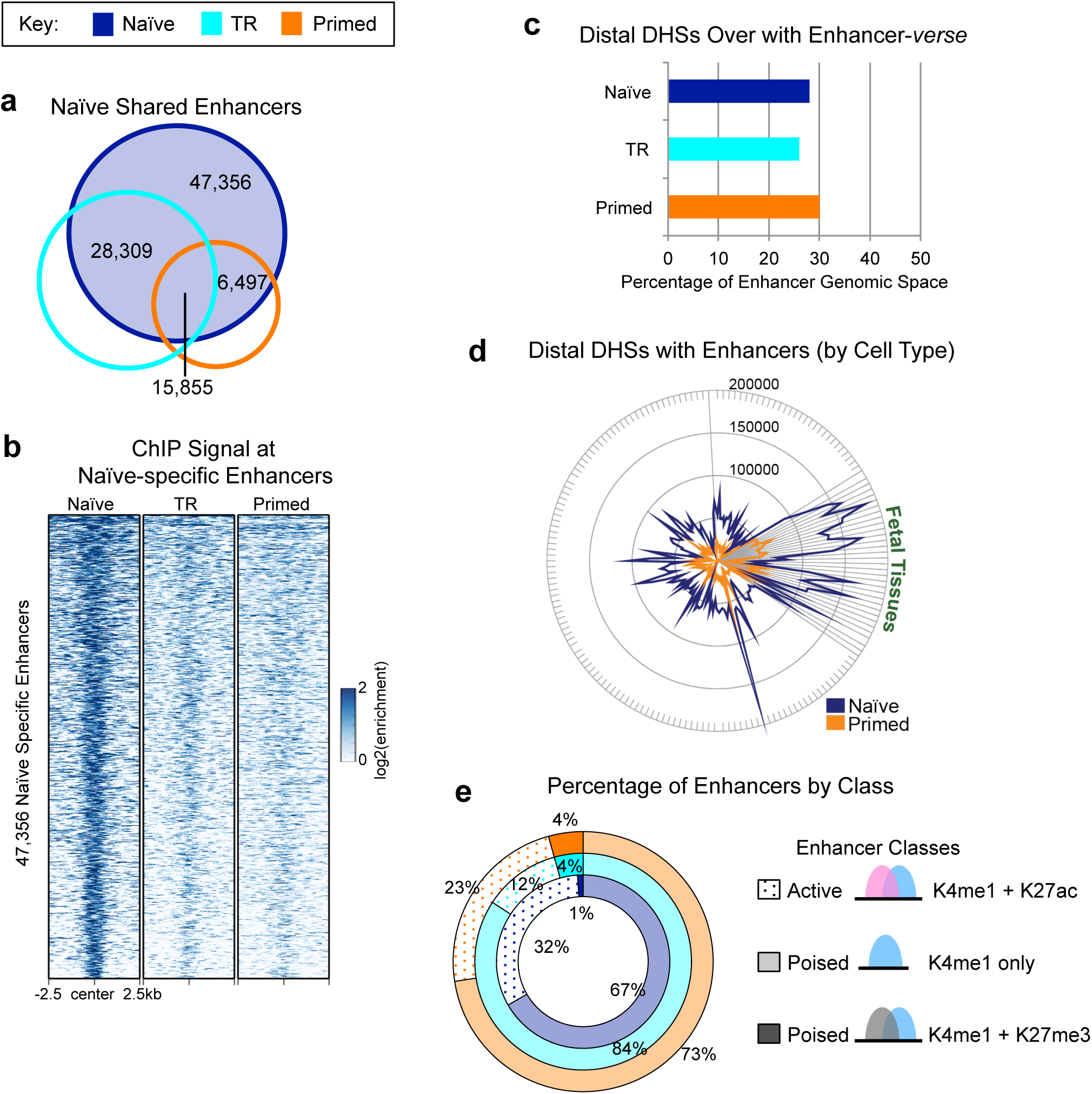
Naïve Enhancer Repertoire. (a) Venn diagram of naïve (navy) enhancers overlapped with transitioning (cyan) and primed (orange) enhancers. (b) Heatmap of H3K4me1 normalized ChIP-seq signal centered at naïve-specific enhancers in a 5kb window. (c) Percent of hESC H3K4me1 genomic space (% bases or enhancer-verse) occupied by ENCODE DHSs from 177 cell types (d) Number of ENCODE DHS from 177 cell types overlapping with naïve and primed H3K4me1 enhancers (e) Distribution of active (H3K4me1 + H3K27ac) and poised (H3K4me1 only or H3K4me1 + H3K27me3) enhancer states in each cell type.

Previous studies, including our own work, in naïve hESCs observed a reduction of H3K27me3 in naïve derived and reset hESCs(Chan et al., 2013; Gafni et al., 2013; Sperber et al., 2015; Theunissen et al., 2014; Ware et al., 2014), consistent with what was shown in naïve mESCs(Marks et al., 2012). Comparisons across cell types reveal a genome-wide depletion of repressive histone modifications in naïve cells (Fig. 2b,c). H3K27me3 repressed regions are more abundant and broader in primed than in naïve cells, covering ∼1.4% of the genome in primed cells compared to 0.5% in naïve (Fig. 2b, Supplemental Figure 3c), which we previously showed is linked to metabolic differences between the cell states(Sperber et al., 2015). H3K9me3 heterochromatin regions, which are sparse in primed cells(Hawkins et al., 2010), are further depleted in transitioning and naïve cells (Fig. 2b,c and Supplementary Figure 3d and Supplementary Table 4,5). There is a notable abundance of H3K4me1 regions in naïve hESCs (Fig. 2b and Supplementary Table 4). Over 9% of the naïve genome is marked by H3K4me1, three times more than primed cells and 1.7 times more than transitioning cells (Fig. 2c and Supplementary Table 5). Monomethylation is present in larger domains, reaching sizes of over 30kb in transitioning cells and over 50kb in naïve cells (Supplemental Fig. 3e). Acetylation is also more enriched in naïve cells with 3x more peaks than primed, and broad H3K27ac domains reaching over 50kb (Fig. 2b,c, Supplemental Figure 3f and Supplementary Table 4,5). The trends for H3K27 modifications also hold true on the X chromosome (Supplementary Fig. 3h-j), where both are active in naïve cells(Ware et al., 2014). We found H3K4me3 to be the most stable mark though cell-specific peaks exist (Fig. 2b,c and Supplementary Fig. 3g).

### Promoter Transitions from Naïve to Primed State

We investigated how DEGs were reflected through promoter chromatin states using >19,000 GENCODE defined autosomal protein coding genes. Over 12,000 promoters are marked with H3K4me3 (Supplementary Figure 4a). We subdivided promoters into six categories: (1) active - H3K4me3 and H3K27ac; (2) poised - H3K4me3 only; (3) bivalent - H3K4me3 and H3K27me3; H3K27ac - H3K27ac only; (5) H3K27me3 - H3K27me3 only; and (6) unmarked - lacking all three modifications (Fig. 2d and Supplementary Figure 4b). Although the largest percentages of gene promoters remain static as either active or unmarked across all three stages, many promoters change chromatin state (Supplementary Figure 4c), which exemplifies the dynamic nature of the epigenome. To illustrate that chromatin patterns coincide with general trends of expression, we plotted the RPKM values of genes with active, poised and bivalent promoters. As expected, genes with active promoters had overall higher expression levels than genes with promoters in the other two categories (Fig. 2e).

**Figure 4.**
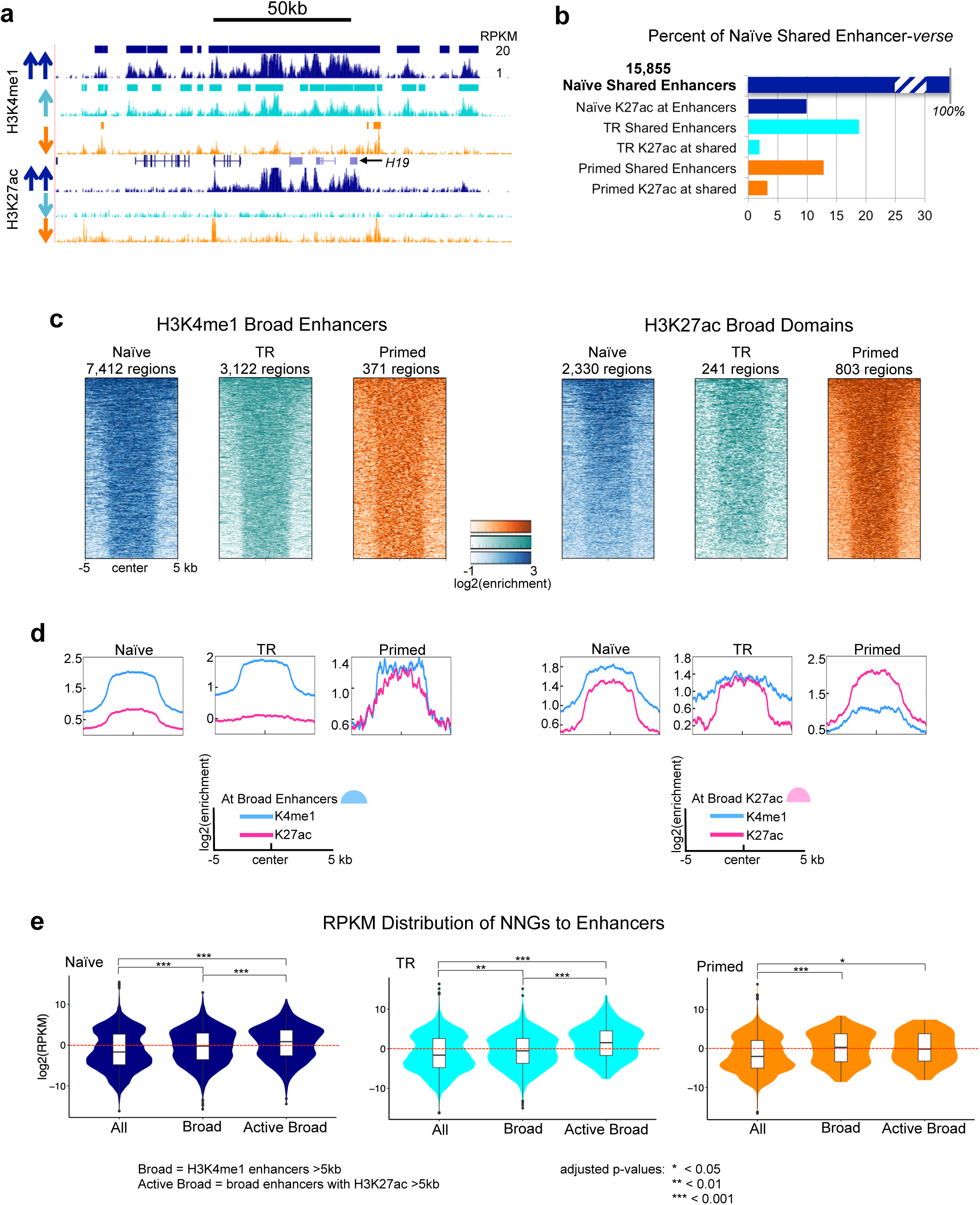
Naïve Enhancers are Decommissioned but Active in Other Cell Types. (a) UCSC Genome Browser image illustrating loss of H3K27ac, followed by loss of H3K4me1 at the *H19* the H19 locus as cells move from naïve (navy), to transitioning (cyan), to primed (orange); RPKM range 1-20 for each track. This region also contains a broad enhancer domain in naïve hESCs. Enhancer peak calls are represented as bars above the H3K4mei track. (b) Percent of shared naïve enhancers genornic space that is preserved in the follow hESC states. © Heatmaps of H3K4mei and H3K27ac normalized ChIP-seq signal at broad enhancer regions (≥5kb). (d) Histograms of average H3K4mei and H3K27ac normalized ChIP-seq signal at all broad enhancers or broad H3K27ac domains. (e) Violin plots showing the distribution of RPKM values of nearest neighboring genes of all broad enhancers (H3K4me 1 ≥5kb) and active broad enhancers (H3K4mel ≥5kb overlapping H3K27ac Z5kb) in each cell type. P-values for pairwise comparisons are computed using two tailed t-tests with pooled SD. P-values are adjusted with Benjarnini-Hochberg method. * P-value < 0.05; ** P-value <0.01; *** P-value <0.001.

Mouse ESCs grown in serum have a greater than three-fold increase in bivalent promoters relative to cells of the mouse ICM (Liu et al., 2016). Observing a similar increase in bivalent gene promoters from naïve to primed cells (1,097 vs 2,674), we determined from which epigenetic states the primed bivalent promoters arose. Roughly 60% of primed bivalent promoters are bivalent in transitioning cells, and of those, their promoter states are split between active (42%), bivalent (32%) and poised (20%) in naïve hESCs (Fig. 2f). Of the ∼7% of naïve active gene promoters that become bivalent in transitioning cells, these genes were enriched for GO terms such as morphogenesis and WNT signaling, and includes genes such as *HOXA1*, *HOXA4*, *HOXD8* and *ZEB1*. Naïve bivalent genes fall into categories involving GO terms for synaptic transmission, ion transport and neuron differentiation (Fig. 2g). Thus, it appears that the neural lineage is the first lineage to be bivalently marked in naïve cells and suggests that naïve hESCs may be an excellent model for further investigation of the establishment of Polycomb repressive regions in the early epigenome.

### Enhancers in the Naïve Embryonic State

Enhancer elements are *cis*-acting regulatory sequences that control gene expression via interaction with transcription factors and promoters. Enhancer chromatin modifications are highly dynamic and cell type-specific(Hawkins et al., 2010). Here, we defined enhancers as H3K4me1 peaks lacking overlap with H3K4me3 (Supplementary Table 6). Investigation of the enhancer landscape across hESC states revealed that naïve cells harbor the most cell type-specific enhancers (>47k; Fig. 3a,b), while transitioning and primed cells had roughly the same number of unique enhancers at ∼17k and ∼14k respectively (Supplementary Figure 5a-d). Sixty-four percent of transitioning enhancers and 55% of primed enhancers are marked in the naïve state (Supplementary Figure 5a,b). We asked if the expansion of naïve H3K4me1 was random or occurred at known regulatory elements. Using DNase I Hypersensitive Sites (DHS) data from 177 ENCODE cells(Consortium, 2012), including H1, we found 25-30% of the H3K4me1-marked genome (enhancer-verse) to be hypersensitive in each cell type (Fig. 3c). Of the 177 cell and tissue types, fetal tissues had the largest collection of DHS overlapping naïve enhancers (Fig. 3d and Supplementary Table 7). Additionally, over 92% of the enhancer base pairs covered by naïve H3K4me1 peaks are utilized as enhancers in 127 Roadmap Epigenome Project cell types, as indicated by H3K4me1 (Supplementary Figure 5e-f). Single cell RNA-seq data from early human embryogenesis(Yan et al., 2013) indicates that 92% of annotated transcription factors(Zhang et al., 2015) are expressed by the late blastocyst stage (Supplemental Figure 5g). Their expression provides a plausible means for aiding the localization of H3K4me1 to known enhancers.

**Figure 5.**
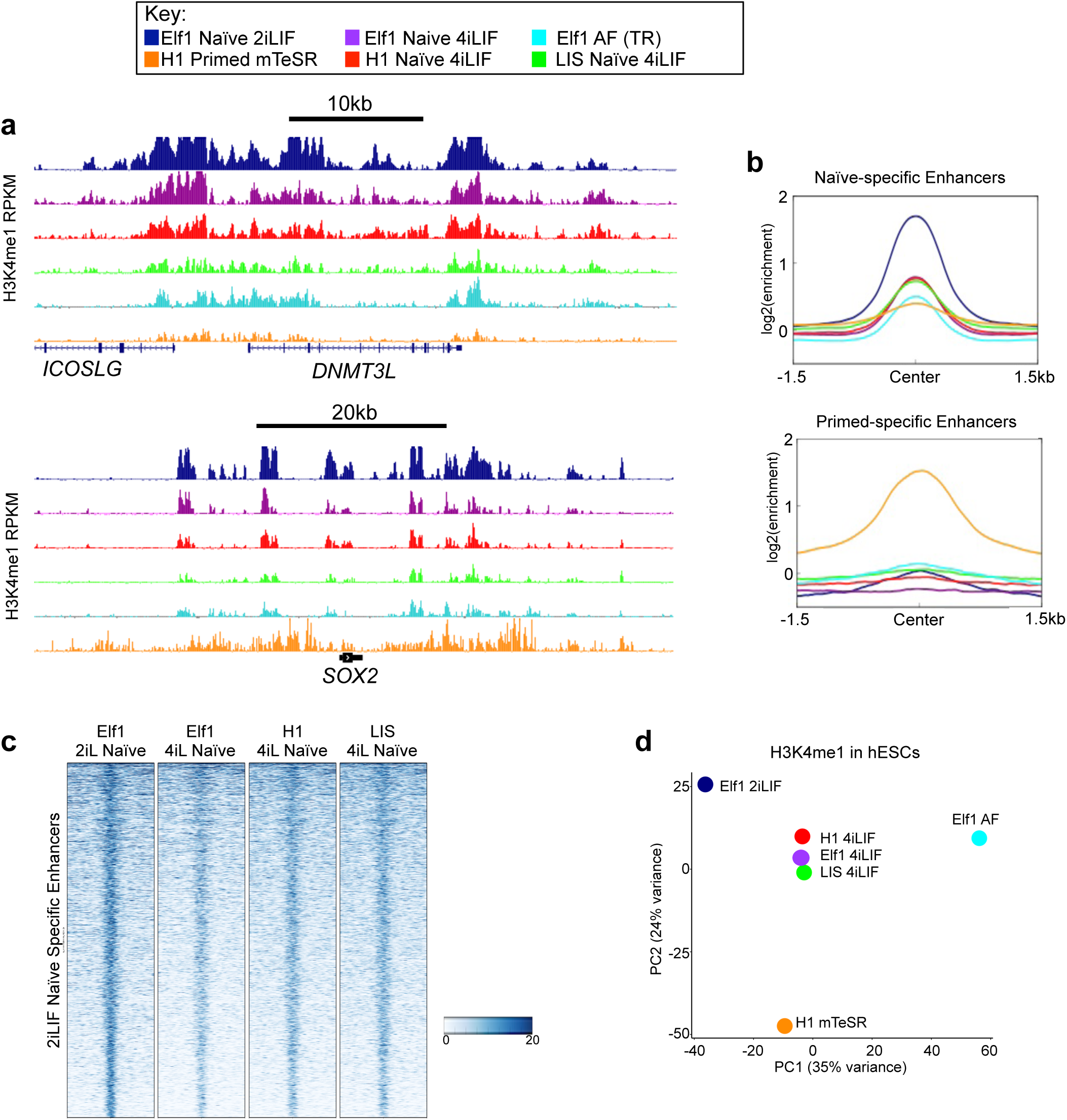
Naïve Enhancers from Various Naïve Culture Conditions. ChIP-Seq of naïve cells grown in different culture conditions including naïve Elf1 naïve (navy), Elf1 4iLIF (purple), Elf1 AF (transitioning - cyan), primed H1 mTeSR (orange), naïve H1 4iLIF (red), naïve LIS1 4iLIF (green). (a) H3K4me1 enrichment in different growth conditions at *DNMT3L* (top panel) and *SOX2* loci (bottom panel) (b) Average ChIP-Seq signal at naïve-specific enhancers (top panel) and primed-specific enhancers (bottom panel). (c) Heatmap of H3K4me1 ChIP-Seq signal at naive specific enhancers. (d) PCA of top 500 10kb bins of H3K4me1 with largest variance.

Enhancer elements can exist in distinct chromatin states that indicate whether they are active or poised (Fig. 3e)(Creyghton et al., 2010; Hawkins et al., 2011; Rada-Iglesias et al., 2011). We characterized differences in the classes of enhancers in each hESC state. We defined active enhancers as regions having H3K4me1 and H3K27ac and poised enhancers as regions with either H3K4me1-only or H3K4me1 and H3K27me3. In all three stages of pluripotency, the majority of enhancers are in the H3K4me1-only poised state (67%, 84%, and 73% in naïve, transitioning and primed cells respectively; Fig. 3e). There is an increase of H3K27me3 containing poised enhancers moving from naïve to primed (1% to 4%; Fig. 3e), which correlates with the increase of H3K27me3.

Our comparative analysis of enhancers indicates that both active and H3K4me1-only poised enhancers are largely decommissioned as naïve hESCs transition to the primed state (Fig. 4a). When assessing overlapping H3K4me1 peaks across hESCs, we see that the chromatin-marked genomic space of naïve enhancers is greatly reduced in primed cells (Fig. 4a,b). This process happens in a stepwise manner, as is evidenced by the loss of acetylation as cells exit the naïve state followed by the gradual loss H3K4me1 (Fig. 4a,b). This introduces a different view of development compared to previous studies that showed poised enhancers gain acetylation following differentiation and were often enriched near genes that became activated later in development(Creyghton et al., 2010; Hawkins et al., 2011; Rada-Iglesias et al., 2011). By using naïve hESCs as a model, we can infer that not only is H3K4me1 likely maintaining open chromatin to aid in the pluripotency phenotype, but that a substantial fraction of enhancers in the human genome are pre-marked early during embryogenesis and subsequently decommissioned during priming.

### Broad Enhancer Domains in the Naïve Epigenome

Super(Whyte et al., 2013) and stretch(Parker et al., 2013) enhancers, which are largely based on H3K27ac, were originally identified in primed ESCs. These regions were shown to upregulate nearby genes and were stronger than conventional enhancers. We asked to what degree these regions were present in our naïve hESCs. To identify both broad H3K4me1 and H3K27ac domains, we identified regions ≥5kb in all cell types (Fig. 4c). The H3K4me1 broad enhancers are almost 20 times more abundant in the naïve epigenome compared to the primed hESC stage (7,412 in naïve hESCs compared to 371 in primed) with an average size of 8.1kb compared to 6.1kb in primed (Supplementary Figure 5h,i). The number of broad enhancers steadily declines as hESCs transition from naïve to primed. We observed the same trend with H3K27ac broad domains (2,330 in naïve compared to 803 in primed), although the number of broad H3K27ac domains in naïve cells is three times less than the number of H3K4me1 broad enhancers (Supplementary Figure 5h). As a control, we looked for broad H3K4me3 peaks, which were limited across the different hESC stages (Supplementary Figure 5h).

Next, we determined if H3K4me1 broad enhancers and H3K27ac broad domains occupy the same genomic space. The average number of bases contained within the overlap of broad H3K4me1 and H3K27ac domains is over 70% of the average length of each domain (Supplementary Figure 5i). Over 78% of broad H3K27ac domains in naïve cells are found within H3K4me1 broad enhancers (Supplementary Figure 5j). In the naïve and primed states 87% and 71% of H3K4me1 broad enhancers, respectively, contained some overlap with H3K27ac, indicating that they are active enhancers (Fig. 4d and Supplementary Figure 5j). The average ChIP-seq signal for H3K4me1 is high at H3K27ac broad domains in all cells except primed hESCs (Fig 4d). The active state of broad enhancers is supported by the distribution of expression values of nearest neighboring genes (NNGs; Fig. 4e). Only in the primed state are there more broad H3K27ac domains than H3K4me1 domains and the difference in the expression distribution of NNGs at broad enhancers versus active broad enhancers in primed cells was the only comparison not found to be significant (Fig. 4e). This may explain why H3K27ac was originally associated with “super/stretch” enhancers. The frequent occurrence of H3K4me1 and H3K27ac broad domains, where broad H3K27ac domains lie within broad enhancers, provides an additional means of giving the genome its “open structure” in naïve pluripotency.

### Naïve hESCs Enhancers in Different Growth Conditions

To determine if the expansion of H3K4me1 in the naïve epigenome was indicative of the naïve state and independent of a single growth condition or cell line, we grew three lines in 4i (2i + p38 kinase inhibitor + JNK inhibitor) + Lif + IGF1 + FGF (referred to as 4iLIF): Elf1, H1 reset to naïve and the naïve derived LIS1 line(Gafni et al., 2013), which grew slightly better in 4iLIF compared to the original growth conditions and compared their transcriptomes for similarity (Supplemental Figure 6a,b). In order to determine the effect of growth conditions and genetic background on the enhancer landscape, we compared the enhancer profiles from H3K4me1 ChIP-seq data across cell types and conditions. Overall, all naïve cells have a similar enhancer profile (Fig. 5a). Cells grown in 4iLIF exhibit a stronger enhancer signal at Elf1 2iLIF naïve-specific enhancers (Fig. 5b,c), and less enrichment at primed- and transitioning-specific enhancers (Fig. 5b and Supplementary Figure 6c). PCA of H3K4me1 signal reveals that all lines grown in 4iLIF are largely indistinguishable, and most similar to 2iLIF (Fig. 5d). Transitioning cells (Elf1 AF) have naïve-like enhancer profiles as mentioned above, transitioning cells have lost naïve H3K27ac but have not yet lost H3K4me1 to primed levels. Our analysis suggests that naïve 2iLIF enhancers do not vary greatly in 4iLIF naïve conditions, although 2iLIF naïve hESCs have some distinct H3K4me1 features. The expansion of H3K4me1 regardless of cell line or growth condition confirms this as a new signature of the naïve hESC state. The acquired expansion upon resetting primed H1 cells to naïve may suggest that this epigenetic feature is necessary for maintenance in the naïve state. Further experiments will be needed to confirm this hypothesis.

**Figure 6.**
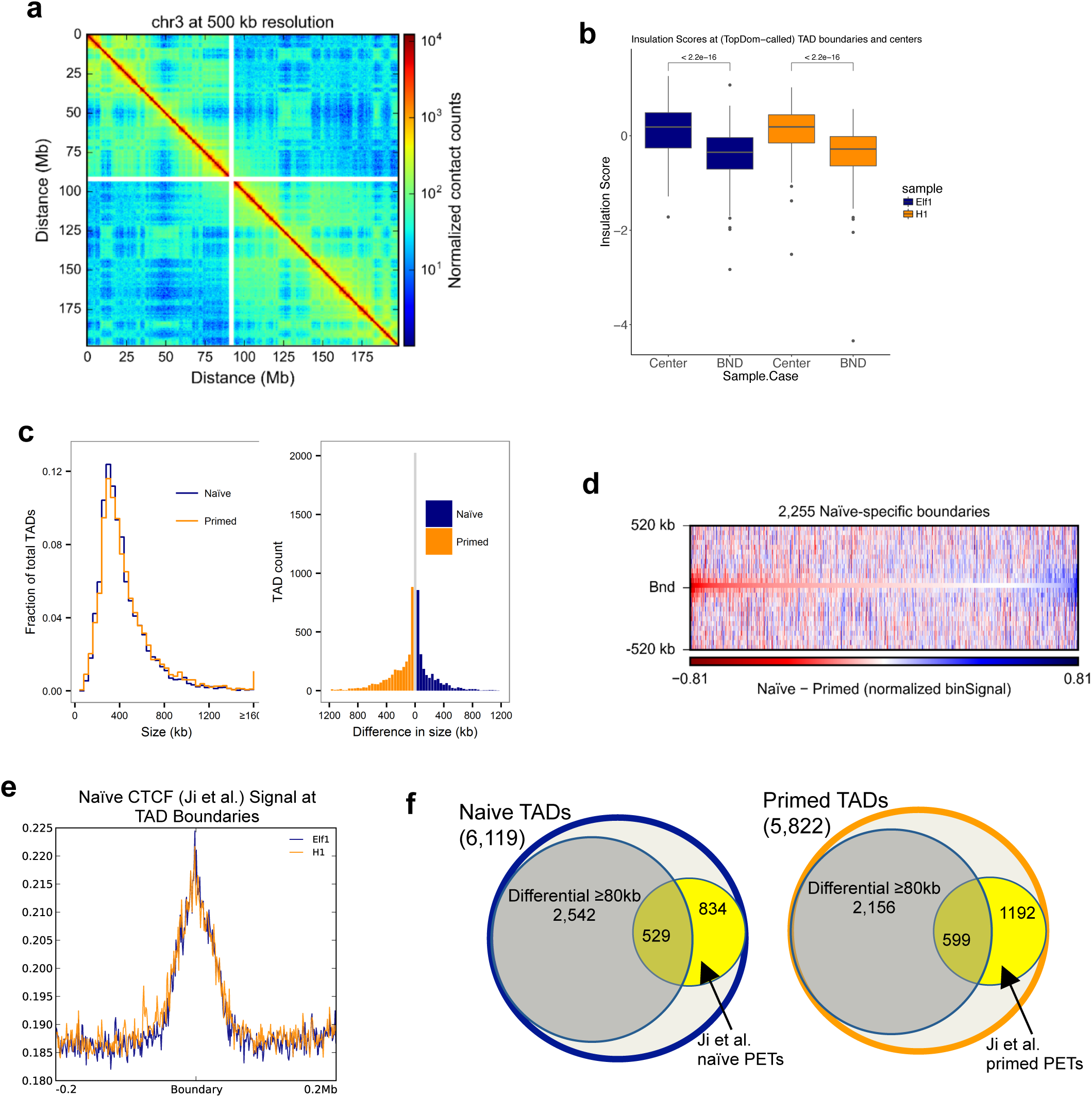
3D Genome Architecture in Naïve hESCs. (a) Hi-C contact heatmap of chromosome 3 in naïve cells at 500kb resolution. (b) Boxplots of the insulation scores along chr7 at both TAD centers and boundaries for both naïve and primed cells. P-values are computed using individual Wilcoxon signed-rank tests. (c) Global size distributions of TADs within naïve and primed cells (left panel) and size differences of overlapping TADs (40kb bin resolution) in naïve and primed cells (right panel). (d) Differential heatmap of naïve minus primed Hi-C bin signal centered at naïve-specific boundary regions. Negative (red) values indicate a stronger bin signal in primed cells relative to naïve cells. (e) Naïve CTCF ChIP-seq signal from Ji et al. 2016, centered at TAD boundaries. (f) Number of TADs or differential TADs with cohesin ChIA-PETs within 40kb of boundary.

### 3D Genome Architecture in Naïve hESCs

Genome architecture is an important component of gene regulation. Topological associated domains (TADs) identified in primed hESCs proved to be surprisingly stable upon differentiation to distinct cell types in spite of diverse changes to chromatin structure(Dixon et al., 2015). Similarly, recent ChIA-PET data for cohesin in primed and naïve reset cells showed a similar recovery of primed TADs(Ji et al., 2016). However, domain-scale 3D genome architecture is still missing for the naïve state. To characterize TADs in naïve Elf1 2iLIF hESCs, we generated deeply sequenced *in situ* DNase Hi-C maps(Deng et al., 2015) (Supplementary Figure 7a), which exhibited characteristic reductions in contact frequency as a function of linear distance between two loci (Fig. 6a). We processed raw Hi-C read pairs produced from H1 primed hESCs(Dixon et al., 2015; Dixon et al., 2012) and compared the architectural features identified In each cell type at 40kb resolution. A total of 6,119 TADs were identified in naïve hESCs compared to 5,822 TADs in primed hESCs (Supplementary Figure 7b), consistent with previous observations in primed hESCs(Shin et al., 2016). We defined boundaries as regions between two adjacent TADs and found that 7.3% and 6.2% of boundaries were greater than 40kb in naïve and primed cells, respectively (Supplementary Figure 7c). To give confidence in our TAD calls, we calculated insulation scores (Crane et al., 2015; Giorgetti et al., 2016). Insulation scores are calculated at each Hi-C bin by aggregating the contact measurements in a fixed window around each Hi-C bin. The insulation score represents how insulated each bin is from TAD boundaries. It is expected that TAD boundaries occur at the valleys/minima of insulation scores, and TAD centers occur at the peaks/maxima. We found that boundary insulation scores were significantly different from TAD center scores (Fig. 6b).

**Figure 7.**
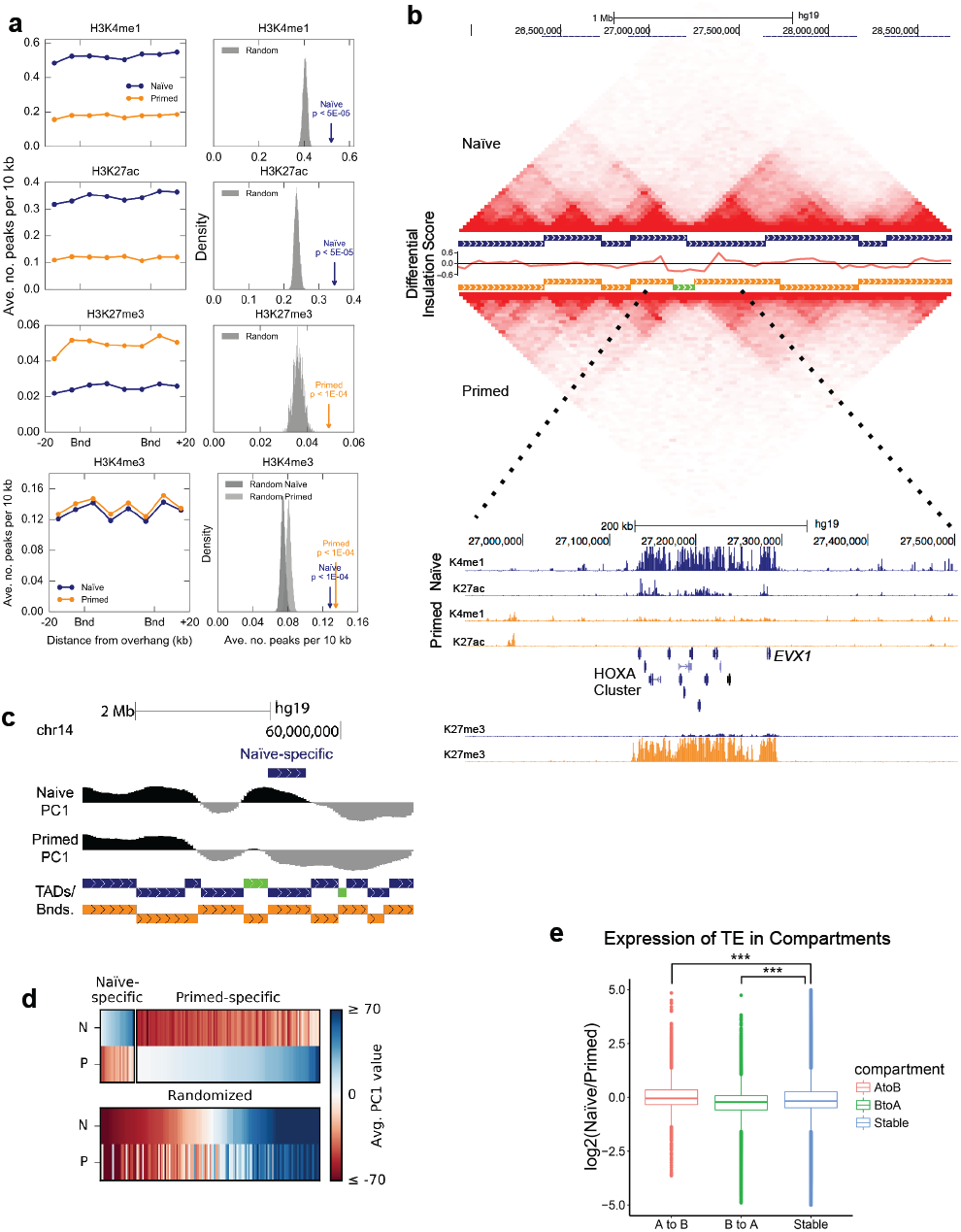
(a) Enrichment of ChIP peaks for histone marks H3K4me1, H3K27ac, H3K27me3 and H3K4me3 at overhanging TAD regions. (b) Interaction matrices of region of chr7 containing HOXA locus. Between matrices, horizontal bars with a vertical offset represents an individual TAD, naïve in navy and primed in orange. The green bar indicates a boundary region > 40kb. Track of differential insulation score of naïve vs. primed cells around the HOXA locus nested in between TAD calls. ChIP-seq signal (RPKM) scaled from 0 to 20 for H3K4me1 and H3K27ac, scaled 0 to 30 for H3K27me3, naïve in navy and primed in orange. (c) Example of a naïve-specific A compartment relative to primed. PC1 scale from −60 to 60. (d) Heatmap of PC1 values at naïve- and primed-specific A compartments. PC1 values at randomized compartments are displayed underneath. “N” and “P” denote naïve and primed, respectively. (e) Boxplot of transposable elements expression (RPKM) overlapping A to B compartment switches. “A to B” and “B to A” are naïve to primed directions. Stable are compartments that do not switch. P-values are computed using two-sample t-test with one sided alternative. *** P-value < 2.2 x 10^−16^.

Overall, TAD size distributions are similar (Fig. 6c, first panel), with means of 420kb in naïve and 444kb in primed. We observed 2,024 TADs whose genomic coordinates are identical at 40kb resolution while the remaining overlapping TADs differ by at least 40kb (Fig. 6c, second panel). We asked if the higher number of naïve Elf1 TADs may be due to better resolution of our *in situ* data, as the two datasets were generated using different Hi-C protocols, and indeed we found that some H1 TADs were split into two or more Elf1 TADs, which accounts for an “extra” 427 naïve TADs (Supplementary Figure 7d). The average overlap between naïve and primed TADs is 319kb, suggesting that the overall TAD structure remains intact between the naïve and primed states (Supplementary Figure 7e). However, we could detect differences in the location of some TAD boundaries as illustrated by naïve-specific boundaries exhibiting an enrichment of primed Hi-C signal (Fig. 6d). That is, in many instances when a change in TAD boundary occurs, this is a shift of the boundary at one end of the TAD relative to the other cell type.

Another group performed cohesin ChIA-PET and CTCF ChIP-seq on primed and reset naïve hESCs(Theunissen et al., 2014) which revealed that looping structures can change between the two cell types(Ji et al., 2016). It was also found that most of the previously published H1 TADs(Dixon et al., 2015) have a CTCF binding site near their boundaries. We asked if the reset naïve CTCF ChIP-seq signal was also enriched at TAD boundaries in our naïve derived hESCs. We found the CTCF signal to be enriched near naïve boundaries; however, this enrichment was also present for primed hESCs CTCF signal (Fig. 6e). This is expected as Ji et al. found that 80% of CTCF binding sites were common between their reset naïve and primed hESC lines(Ji et al., 2016). This helps confirm our *in situ* DNaseI Hi-C data as accurately capturing the 3D structure of the naïve genome.

Cohesin ChIA-PET data from primed and reset naïve hESCs could recapitulate Hi-C TADs(Ji et al., 2016). Although the authors note that their ChIA-PET data were undersaturated,we asked if the cohesin PETs could help to validate our TAD calls. An overlap analysis with cohesin ChIA-PET data yielded 1,363 naïve and 1,818 primed TADs with at least one PET whose termini are located within 40 kb of each boundary of a given TAD (Fig. 6f). This corresponds to 22% of our naïve TADs having a naïve PET and 31% of primed TADs having a primed PET within 40kb of the TAD boundary. We looked to see if any of the PETs were near (within 40kb) differential TAD boundaries, those having different boundaries in naïve and primed of 80kb or greater. Of 1,363 PETs near a naïve boundary, 529 (39%) are near a naïve differential TAD (Fig. 6f). This helps confirm some of the structural differences observed in the naïve 3D genome.

We investigated if there was a relationship between higher-order chromatin structure at differential TAD boundaries and changes in chromatin modifications. We observe a significant enrichment for H3K4me1 and H3K27ac at the internal edge of TADs with differential TAD boundaries in the naïve state relative to random (naïve H3K4me1 and H3K27ac P-value < 5x10^-5^), and a similar enrichment for primed H3K27me3 (P-value < 1x10^-4^) (Fig. 7a). A clear example illustrating these differences in TAD and chromatin structure is the HOXA cluster, where a broad boundary spans the HOXA cluster in primed hESCs and is enriched for H3K27me3 (Fig. 7b). In naïve hESCs, where HOXA genes are expressed, the TAD to the left of the boundary in primed cells is extended across the cluster and marked by H3K4me1 and H3K27ac. We calculated the differential insulation scores by comparing the naïve minus primed insulation scores to confirm a significant difference in the TAD structures. The differential insulation score represents the differential TAD structure between two samples. We examined the differential insulation score around the HOXA locus (Fig. 7b), and observed that there was a noticeable decrease in the signal at the HOXA locus, confirming that the TAD structure at the HOXA locus is different between naïve and primed cells.

Finally, to compare the spatial organization of chromatin within the nuclei of naïve and primed hESCs, we partitioned the genome into active and inactive (A/B) compartments by performing a PCA of each intra-chromosomal contact matrix(Dixon et al., 2015; Lieberman-Aiden et al., 2009). Compartments identified using the first principal component (PC1) ranged in size from 40 kb to over 49 Mb in both cell types, with means of 3.6 Mb in naïve cells and 3.4 Mb in primed. An overwhelming majority of compartments are static, with only 23 switching from being active in naïve cells to inactive in primed (A to B), and 124 switching from being active in primed cells to inactive in naïve (B to A; Fig. 7c,d). While there is enrichment of primed-specific active compartments, a previous study showed that inactive B sub-compartments are largely devoid of histone modifications, including H3K27me3 and H3K9me3(Rao et al., 2014). It is therefore likely that the primed-specific active compartments are driven by the lack of repressive modifications in naïve hESCs (alternatively, these are naïve-specific inactive B compartments). Additionally, cell-specific active compartments are enriched for TE expression relative to stable compartments (Fig. 7e), and gene expression to a lesser extent (Supplementary Figure 7f).

## DISCUSSION

Embryogenesis represents the most dynamic epigenetic reprogramming event in mammalian biology. The best illustrative example of this is the erasure of DNA methylation after fertilization that is eventually reestablished upon implantation to prepare the embryo for further development. Coinciding chromatin dynamics upon implantation are largely still unknown. The best supporting evidence that naïve cells are representative of the pre-implantation embryo is their DNA hypomethylation state, and the corresponding expression of non-coding RNAs such as ERVs and TEs (Grow et al., 2015; Theunissen et al., 2016). Thus, naïve and primed ESCs provide a system to model the transition from the pre- to post-implantation state of the embryo and reveal additional epigenetic changes.

The epigenetic states of embryonic development are also fundamental to our understanding of pluripotency and developmental competency of ESCs. The leading hypothesis for how pluripotency is conferred in ESCs is through a more open chromatin structure relative to other somatic cells, including relative to other stem or progenitor cells (Hawkins et al., 2010; Hiratani et al., 2010; Meshorer et al., 2006). This open chromatin state must, therefore, be derived earlier during embryogenesis. Various new ESC culture conditions now capture different points along the spectrum of pluripotency. Many of which may reflect embryonic stages of development. Our comprehensive chromatin state analyses provide further insight on how naïve cells are unique from primed hESCs, and exist in a distinctly open chromatin state. A number of recent studies on the genome-wide localization of histone modifications, as well as 3D genome architecture, during mouse embryogenesis provide a framework for a contextual understanding of open chromatin in naïve hESCs.

During mouse embryogenesis, promoter H3K27me3 accumulates during embryogenesis and implantation (Liu et al., 2016; Zheng et al., 2016). The cells of the blastocyst ICM exhibit reduced H3K27me3 as do naïve (2i-Lif) mESC, while meta-stable mESCs (Lif+serum) exhibit an increase of H3K27me3 relative to naïve mESCs (Zheng et al., 2016). The depletion of H3K27me3 and far fewer bivalent promoter chromatin states in naïve hESCs relative to primed cells is consistent with this pre-implantation chromatin signature. We determined 1097 bivalent genes in naïve hESCs. This is consistent with mouse morula and ICM embryo stages where approximately 1000 and 2000 bivalent gene promoters exist, respectively (Liu et al., 2016). Another striking similarity is that naïve bivalent genes are enriched for GO terms related to neurogenesis, which is also true for genes gaining bivalency leading up to mouse ICM formation (Liu et al., 2016).

Within hours of fertilization of the mouse oocyte, H3K4me1 begins to increase. This initially begins on the paternal genome around five hours post fertilization (p.f.) (Lepikhov and Walter, 2004), and just after incorporation of histone H3 (van der Heijden et al., 2005). Increased H3K4me1 coincides with the period of active DNA demethylation of paternal DNA (Santos et al., 2002), after which H3K4me1 continues to increase on both genomes. Our data show that H3K4me1 has undergone massive expansion in naïve hESCs relative to primed cells. Although expanded H3K4me1 regions mark known human enhancers, they exist in a poised enhancer chromatin state (H3K4me1), and are decommission upon transition to the primed state, rather than being poised for activation at the next developmental stage.

Enhancer decommissioning, through LSD1 activity, is required for proper ESC differentiation (Whyte et al., 2012). LSD1 activity is inhibited by acetylation(Forneris et al., 2005; Lee et al., 2006) which suggests that de-acetylation must precede the removal H3K4me1 by LSD1. We observed this stepwise decommissioning as cells exited the naïve state. Recently, ChIP-seq results for H3K27ac in the mouse embryo and serum-maintained mESCs showed enrichment of H3K27ac genome-wide post-ZGA followed by a decline in mESCs(Dahl et al., 2016). This lends support to our hypothesis that enhancer pre-marking is a likely component of epigenetic reprogramming during embryogenesis.

Because most human TFs are expressed during embryogenesis (Yan et al., 2013), one hypothesis would be that the increased abundance of H3K4me1 could be the remnants of once active enhancers from earlier in embryogenesis. However, upon resetting primed H1 hESCs to a naïve state, we found that the expansion of H3K4me1 was gained. This indicates that expansion H3K4me1 domains are a hallmark of the naïve, pre-implantation state and may have an alternative function.

Because expansion of H3K4me1 across the genome coincides with the hypomethylated DNA state of both naïve hESCs and the early mouse zygote, we posit that the expansion is either necessary for or aids in maintaining DNA hypomethylation. Although H3K4me3 is primarily cited as the modification being mutually exclusive to DNA methylation, all methylation states of H3K4 (mono-, di-, and tri-) were originally shown to inhibit binding of DNMT3L to the histone H3 tail (Ooi et al., 2007), which upon binding recruits the *de novo* methyltransferases DNMT3A and 3B. Somewhat paradoxically, our expression data, and that of others, shows that *DNMT3L* is upregulated in naïve hESCs (Blakeley et al., 2015; Sperber et al., 2015). Future studies are needed to understand the interplay between expanding H3K4me1 and DNA demethylation in the pre-implantation state.

Furthermore, the additional depletion of H3K9me3 in the naïve state signifies an overall dramatic reduction in epigenetic repression relative to primed cells: reduced DNA methylation, H3K27me3 and H3K9me3. Similarly, H3K9me2 is rapidly removed from the maternal genome of the mouse embryo shortly after fertilization (Lepikhov and Walter, 2004; Sarmento et al., 2004; van der Heijden et al., 2005). This is accompanied by hyperacetylation of histones (Adenot et al., 1997; Wiekowski et al., 1997). Our data also show an increase in histone acetylation in naïve state relative to primed. Collectively, these events are likely important in establishing an open chromatin state in the embryo and ESCs.

Naïve versus primed hESCs illustrate one of the more dramatic changes in chromatin architecture shown between two cell types. We asked if these changes impacted 3D genome structure by generating HiC interaction maps in naïve hESCs. We found that over 2000 TAD boundaries shifted by at least 80kb between the cell types. As confirmation that the changes reflect biological differences, we found that naïve-specific boundaries overlap recently published CTCF binding sites from hESCs reset to the naïve state. However, these sites are also bound by CTCF in primed hESCs, which may suggest the boundaries shift from one CTCF site to another and other factors may control the boundary position. Consistent with this, overlap of recent cohesin ChIA-PET data revealed that of PETs at naïve TAD boundaries, 39% were localized to naïve-specific boundaries. TAD structures are reported to be stable across cell or tissue type, yet these comparisons also reveal specific TAD boundaries (Dixon et al., 2012; Schmitt et al., 2016). Some boundary differences are, therefore, likely expected in any pairwise comparison. In addition, two recent reports showed a gradual, step-wise establishment of TADs during pre-implantation embryogenesis (Du et al., 2017; Ke et al., 2017). A comparison of TADs from the mouse ICM to meta-stable mESCs (grown in serum plus Lif) showed 80% overlap in TAD boundaries (Du et al., 2017). Although meta-stable mESCs exist in a state between naïve and primed, the results suggest that TAD finalization continues into the implantation stage of development. As proof of this, mouse E3.5 and E7.5 embryos continued to show TAD rearrangement relative to each other and previous developmental stages (Ke et al., 2017). Therefore as in mouse embryogenesis, TAD differences between naïve and primed hESCs may reflect the continuum of TAD establishment.

In conclusion, naïve hESCs provide an in vitro model system for studying epigenomic dynamics as cells transition to from the pre- to post-implantation state. This system provides novel insight on this highly dynamic event, and suggest hypotheses that can be tested in mouse, or possibly human, embryos. Furthermore, naïve cells reveal a more open chromatin state that is likely reflective of earlier epigenomic events during embryogenesis. Human ESC culture conditions provide a tractable system for further investigation of the interplay between epigenomic modifications.

## MATERIALS AND METHODS

### Human Embryonic Stem Cell Culture

All human ESC culture conditions were as previously described(Sperber et al., 2015), with the following modifications. Growth conditions: 2iLIF - 1uM Mek inhibitor (PD0325901) [catalog #S1036, Selleck Chemicals, Houston, TX, USA], 1uM GSK3 inhibitor (CHIR-99021) [catalog #S2924, Selleck Chemicals, Houston, TX, USA], 10 ng/mL Leukemia inhibitory factor [catalog #YSP1249, Speed Biosystems, Gaithersburg, MD, USA], 5ng/mL IGF-1 [catalog #100-11 Peprotech, Rocky Hill, NJ], 10ng/mL FGF [catalog #PHG0263, Thermo Fisher Scientific, Waltham, MA, USA]; 4iLIF - 1uM Mek inhibitor (PD0325901), 1uM GSK3 inhibitor (CHIR-99021), 5uM JNK inhibitor (SP600125) [catalog #S1460, Selleck Chemicals, Houston, TX, USA], 2uMp38 inhibitor (BIRB796) [catalog #S1574, Selleck Chemicals, Houston, TX, USA], 10 ng/mL Leukemia inhibitory factor, 5ng/mL IGF-1, 10ng/mL FGF.

### Chromatin Immunoprecipitation and Sequencing (ChIP-seq)

ChIP-seq was performed as previously described(Hawkins et al., 2013). Raw sequence reads from Roadmap Epigenome Project(Hawkins et al., 2011). All sequenced reads were analyzed with the same pipeline and settings. Sequence reads were aligned to genome (version hg19) using Bowtie2(Langmead and Salzberg, 2012). Replicates of aligned files were merged prior to peak calling. For the UCSC genome browser tracks, ChIP-seq signals were normalized by RPKM followed by subtraction of input from ChIP using deepTools suite(Ramirez et al., 2014). Heatmaps and histograms are of normalized ChIP-seq signal: samples are normalized by read count and log2(chip reads/input reads) per 10kb bin is plotted using deepTools suite(Ramirez et al., 2014).

### Peak Calling

ChIP-seq peaks were called on merged replicates and normalized to input using MACS v1.4(Zhang et al., 2008). Peak calls with a FDR of 5% or less were used for downstream analysis. Percent of genome covered was defined as total number of bases under the peak divided by 2.7e9, the effective genome size. This was found it to be a better representation of global chromatin structure (e.g. a 10kb region can be covered by one or many ChIP-seq peaks due to peak size; the number of peaks may vary more than the total number of bases under the peaks). Peak comparisons and overlaps were done using the BedTools suite(Quinlan and Hall, 2010).

In order to compare the histone marks (H3K4me1 and H3K27ac) across cell types, we divided the genome into 10 kb bins and counted the reads across these 10 kb genomic regions using *featurecounts* in Rsubread package(Liao et al., 2014). Then, PCA was performed on regularized log transformed read count data obtained using DESeq2(Love et al., 2014).

### RNA-seq and Gene Expression

Embryonic stem cells were counted and 200,000 cells were pelleted for RNA extraction using the Qiagen All Prep Kit (cat 80004). RNA-seq libraries were constructed using the Scriptseq RNA-seq Library Preparation Kit on ¾ of total RNA. Libraries were sequenced single-end 75 on Illumina NextSeq. The quality of the reads and contamination of adapter sequences were checked with FastQC tool (http://www.bioinformatics.babraham.ac.uk/projects/fastqc/). Reads were mapped to human hg19 genome (UCSC) using TopHat2(Kim et al., 2013). Transcript quantification was performed by Cufflinks(Trapnell et al., 2010) using GENCODE's comprehensive gene annotation release 19 as reference annotation.

### Differential Gene Expression Analysis

The raw read counts were calculated using *featurecounts* in Rsubread package(Liao et al., 2014) and GENCODE’s release 19 as reference annotation. Differential gene expression analysis was performed with DESeq2(Love et al., 2014) using read counts matrix. Two sets of differentially expressed genes (DEGs) are identified with P-value < 0.01, log2FC > |1| and P-value < 0.01, log2FC > |2|. The P-values were adjusted for multiple hypothesis correction. DEGs in all pairwise sample comparisons were identified. PCA was performed on regularized log transformed read count data from autosomes of top 500 highly variant genes obtained using DESeq2(Love et al., 2014) and plot was generated using ggplot2 in R(Wickham, 2009). For transposable elements (TE) analysis, transcripts were quantified using hg19 UCSC RepeatMasker TE annotation. We considered unique reads as well as multi mapped reads during quantification of TE transcripts. PCA was performed on regularized log transformed read count data of top 500 highly variant TE transcripts obtained using DESeq2.

### Identification of Overrepresented GO Terms and Enriched Pathways

ClueGO(Bindea et al., 2009) was used to identify the overrepresented GO terms and enriched pathways with the data from gene ontology consortium and KEGG pathways database. The input gene lists to the ClueGO were DEGs with P-value < 0.01, log2FC > |1|. We used all genes in the genome as background. The statistically significant GO terms and pathways were filtered with P-value < 0.05 and GO term/pathway should contain at least 5 DEGs. P-values were adjusted with Benjamini Hochberg method for multiple hypothesis correction.

### Sankey Plot

We looked at promoter chromatin state transitions from naïve to primed to gain insight into the establishment of bivalency and other chromatin state changes occurring at gene promoters. In order to accomplish this goal we focused on the over 19,000 autosomal protein-coding gene TSS annotated by GENCODE. We assigned a promoter to a gene if the H3K4me3 peak was within −2kb to +500bp of the TSS. Sankey plot is limited by by the presence of multiple promoters. Sankey plot were created using Google Charts (https://developers.google.com/chart/interactive/docs/gallery/sankey)

### In situ DNase Hi-C

Samples were prepared in a manner similar to Deng *et al.*, 2015(Deng et al., 2015). Briefly, nuclei from ∼5 x 10^6^ cross-linked Elf1 cells were isolated and permeabilized, and chromatin was digested with 4 U DNase I at room temperature for 4 min. Following end-repair and dA-tailing reactions, chromatin ends were ligated to biotinylated bridge adapters, and nuclei were purified with two volumes of AMPure XP beads (Beckman Coulter). Chromatin ends were phosphorylated and ligated in situ, and protein-DNA cross-links were reversed by proteinase K digestion and incubation at 60°C overnight. Following purification, DNA was sonicated to an average size of 400 bp, and chimeric species were enriched via pull-down with streptavidin-coated magnetic beads (Active Motif). Preparation of Hi-C libraries was accomplished by ligating sequencing adapters to the ends of bead-bound DNA fragments and PCR-amplifying the products in the presence of forward and barcoded reverse primers. Libraries were purified with AMPure XP beads, DNA concentrations were determined using a Qubit 2.0 (Thermo Fisher), and size distributions were quantified using a Bioanalyzer with a high sensitivity kit (Agilent). A 10 ng aliquot from each library was digested with BamHI, run on the Bioanalyzer, and compared to an undigested control in order to confirm the presence of a reconstituted BamHI site at the junctions of ligated bridge adapters.

### Hi-C Sequencing and Data Processing

Raw Hi-C sequencing reads from Hi hESCs were downloaded from GEO (GSE35156). Reads were aligned using Bowtie2(Langmead and Salzberg, 2012) to the hgl9 reference genome and filtered for MAPQ ≥ 10, uninformative ligation products, and POR duplicates using HiC-Pro.

Valid Hi-C read pairs from biological replicates of Elf1 and H1 hESCs were combined, respectively, and used to generate raw chromosome-wide interaction matrices binned at a resolution of 40kb. Raw matrices were ICE-normalized using the HiTC Bioconductor package(Servant et al., 2012) for R, and TADs and boundaries were identified using TopDom(Shin et al., 2016) with a window size of 5. X and Y chromosomes were removed for the datasets for all Hi-C analyses.

Insulation scores were calculated for the whole of chr7 from the ICE-normalized matrices of both the Elf1 and H1 hESCs, separately. Insulation vectors were detected via cworld(Giorgetti et al., 2016) using the script matrix2insulation.pl, and using the following options: (--is 240000 -- nt 0.1 --ids 160000 --im median --bmoe 0). Differential insulation scores computing Elf1 score minus H1 score for the whole of chr7 via cworld using the script compareInsulation.pl, with inputs being the two insulation scores above.

High-Confidence SMC1 ChIA-PET interactions for naïve and primed hESCs were downloaded as a supplemental table(Ji et al., 2016). A ChIA-PET was considered to span a TAD if both PET termini were located within 40 kb of a TAD boundary.

Spatial compartments and activity status were identified via principal component analysis (PCA) using Homer Tools(Heinz et al., 2010). Processed Hi-C reads were imported into Homer. For each chromosome, a contact matrix was constructed at 40 kb resolution and normalized using a sliding window of 400 kb as background. Next, the correlation between intra-chromosomal contact profiles was computed and the first principal component (PC1) vector was extracted and saved as a bedGraph file. H3K27ac ChIP-seq peaks served as a seed for determining which regions are active (PC1 > 0). A genomic region was considered cell type-specific if it met the following three criteria: 1) the average PC1 value was positive in one cell type and negative in the other, 2) the difference in the average PC1 value was > 50 and 3) the correlation between contact profiles was < 0.4. Randomization was achieved by selecting coordinates from a pool of 40 kb regions that had associated PC1 values and were not located within any cell type-specific sub-compartments.

## ACCESSION NUMBERS

All data will be submitted to Sequence Read Archive (SRA) upon acceptance.

## SUPPLEMENTAL INFORMATION

Supplementary Information includes 7 figures, and 7 tables.

## ACKNOWLEDGEMENTS

Dr. Gurkan Yardimci (Ph.D., Department of Genome Sciences, University of Washington, Seattle, WA, USA) for communication and advice on Hi-C analysis.

## AUTHOR CONTRIBUTIONS

**COMPETING FINANCIAL INTERESTS**

**MATERIALS & CORRESPONDENCE**

